# ProAffinity-GNN: A Novel Approach to Structure-based Protein-Protein Binding Affinity Prediction via a Curated Dataset and Graph Neural Networks

**DOI:** 10.1101/2024.03.14.584935

**Authors:** Zhiyuan Zhou, Yueming Yin, Hao Han, Yiping Jia, Jun Hong Koh, Adams Wai-Kin Kong, Yuguang Mu

## Abstract

Protein-protein interactions (PPIs) are crucial for understanding biological processes and disease mechanisms, contributing significantly to advances in protein engineering and drug discovery. The accurate determination of binding affinities, essential for decoding PPIs, faces challenges due to the substantial time and financial costs involved in experimental and theoretical methods. This situation underscores the urgent need for more effective and precise methodologies for predicting binding affinity. Despite the abundance of research on PPI modeling, the field of quantitative binding affinity prediction remains underexplored, mainly due to a lack of comprehensive data.

This study seeks to address these needs by manually curating pairwise interaction labels on all available 3D structures of proteins complexes, with experimentally determined binding affinities, creating the largest dataset for structure-based pairwise protein interaction with binding affinity to date. Subsequently, we introduce “ProAffinity-GNN”, a novel deep learning framework using protein language model and graph neural network (GNN) to improve the accuracy of prediction of structure-based protein-protein binding affinities. The evaluation results across several benchmark test sets demonstrate that ProAffinity-GNN not only outperforms existing models in terms of accuracy but also shows strong generalization capabilities.

## Introduction

Protein-protein interactions (PPIs) serve as the cornerstone of nearly all biological processes, dictating the dynamics of signaling pathways and structural frameworks essential for cellular functionality. ^1^ These interactions are central to dissecting the molecular basis of diseases, thus forming a critical focus in the quest for novel therapeutic strategies. ^2,3^ The proliferation of advanced experimental techniques, including X-ray crystallography, NMR spectroscopy, and cryo-electron microscopy, has significantly expanded the repository of 3D structural data on PPIs. This wealth of structural information has set the stage for the integration of artificial intelligence (AI) methods,^4–12^ potentially providing a cost-effective and efficient alternative to traditional experimental approaches.

Beyond the structural delineation of PPIs, binding affinity emerges as a critical determinant, dictating the formation and specificity of protein complexes. ^13^ For instance, this parameter is particularly crucial in drug optimization phases of therapeutic development.^14,15^ Nonetheless, the accurate prediction of protein-protein binding affinities remains a formidable challenge. While traditional experimental methods for affinity determination are known for their resource intensity,^16^ computational approaches, such as molecular dynamics simulations and empirical energy functions, face hurdles in computational demand and accuracy. ^17–22^

In the realm of machine learning, recent endeavors have shown promise for structure-based protein-protein binding affinity prediction.^23–27^ Notably, the PRODIGY predictor harnesses a linear model focusing on inter-residue contacts, non-interacting surface, and buried surface area which are some elements crucial to PPIs.^24^ Similarly, PPI-Affinity employs a Support Vector Machine (SVM) strategy, leveraging selected molecular descriptors as input features.^28^ Despite these advancements, the complex nature of PPIs and the scarcity of comprehensive datasets have hampered the full realization of machine learning’s potential in this field.^29^

PDBbind protein-protein complexes database (version 2020), ^30^ subsequently referred to as PDBbind, is the current largest available PPI structure-based resource, with experimental binding affinities. To map the binding affinity onto the structure features, usually two-body or two-chain of interacting proteins are considered. PDBbind hosts 2,852 protein complexes, however, only 590 of them contain only two chains of proteins which is easy for data preparation, most of the data points contain more than two chains of proteins. This complexity significantly diminishes the original dataset’s applicability for detailed protein-protein binding analysis. In contrast, a dedicated structure-based benchmark database for protein-protein binding affinity,^31^ which can offer a granular view by clearly delineating chain components within pairwise interactions, has only 144 protein-protein pairs, highly limited the potential for comprehensive analysis and development within this research topic.

In this work, our first endeavor is to manually curate each PPI complex in the structure database to identify the interacting two chains or two groups of proteins based on the detailed structural features and related published papers. Leveraging the comprehensive coverage of PDBbind and the precise focus of the structure-based benchmark, we have created a refined dataset with 2,285 unique protein-protein pairs with their binding affinity. The protein-protein pairs in this dataset indicate the components involved in pairwise inter-actions, providing an accurate and extensive foundation for detailed structure-based bench-marking of protein-protein binding affinities prediction. Further details on the construction of this dataset will be elaborated upon in the subsequent sections of this paper.

Building upon our refined dataset, we introduce “ProAffinity-GNN”, an innovative deep learning framework designed to advance the modeling of protein-protein complexes. Leveraging the fusion of protein language model^32,33^ and graph neural networks (GNNs),^34^ ProAffinity-GNN encapsulates both structural and sequence information within residue-level graphs, offering a comprehensive portrayal of protein-protein interactions. This model also significantly enhances performance by collaboratively combining intra- and inter-molecular graphs, providing a detailed and holistic view of protein-protein complexes. This approach enables deeper insights and more precise predictions of binding affinities.

ProAffinity-GNN represents a significant advancement beyond traditional machine learning models for structure-based protein-protein binding affinity prediction, which typically focused on two-chain interactions and relied on the extraction of complex physicochemical properties.^23,27,28^ To our best knowledge, this is the first purely deep learning-based approach for the structure-based prediction of protein-protein binding affinities. ProAffinity-GNN has shown enhanced accuracy and extensive generalizability across diverse benchmark test sets. This work contributes to the field of protein-protein binding affinity prediction, providing insights that may inform future research in protein design and therapeutic target exploration.

## Materials and methods

In this section, we first introduce the details of the construction of our protein-protein binding affinity dataset. Then, we elucidate the proposed ProAffinity-GNN (see Figure 1), including data processing, graph construction, and network training.

**Figure 1:**
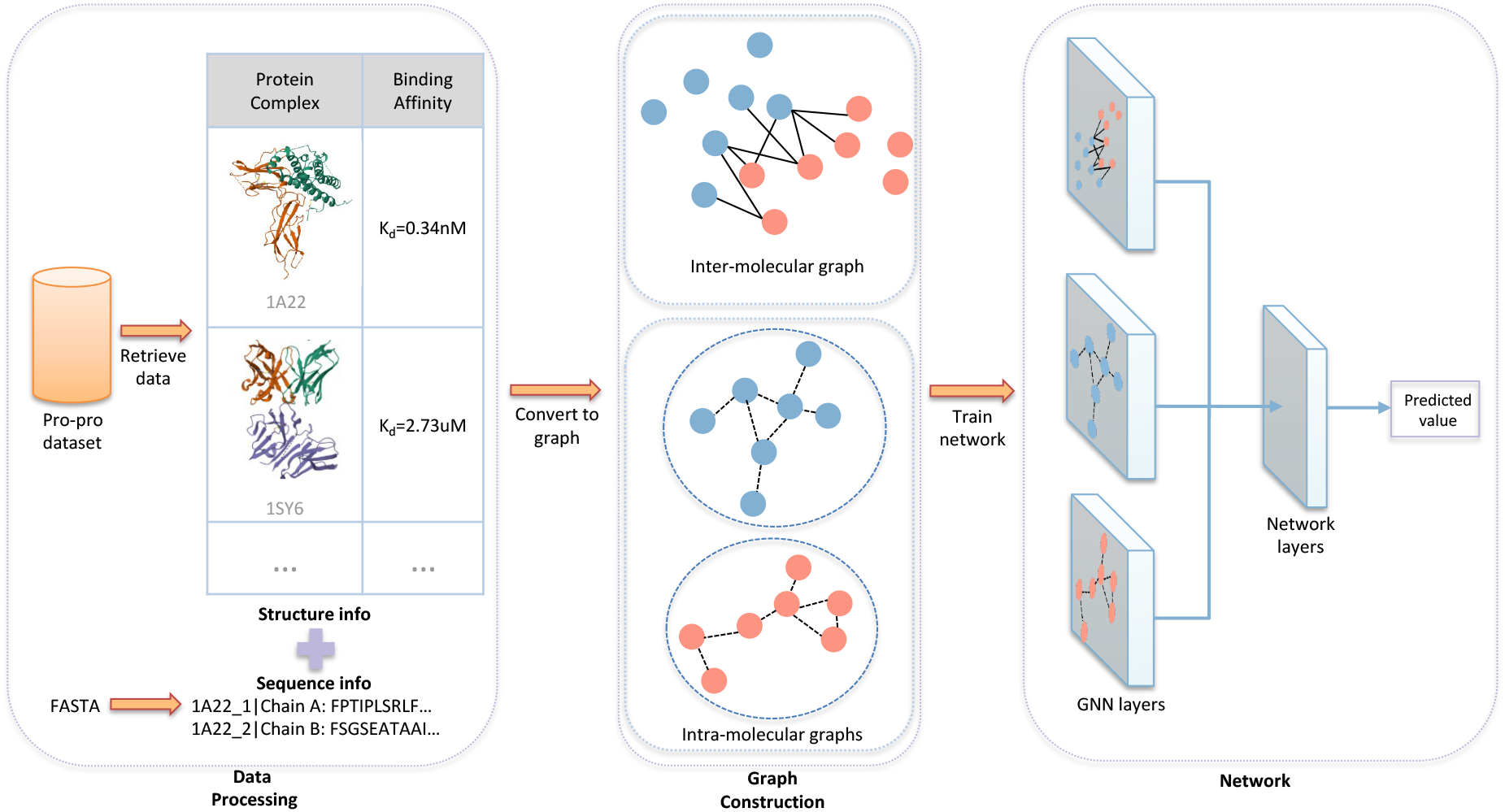
Pipeline of ProAffinity-GNN. First, 3D structure and sequence data of the protein-protein complexes with corresponding binding affinities are retrieved. Then, one intermolecular and two intra-molecular graphs are constructed based on the data. Finally, a network is trained, with the graphs as the input.

### Dataset creation: structure-based protein-protein binding affinity

In this study, we constructed a dataset for structure-based protein-protein binding affinity prediction, utilizing the PDBbind^30^ database as a foundation. The original PDBbind dataset contains 2,852 entries, each comprising a 3D structure in PDB format and experimentally determined binding affinity values, represented as *K*_*d*_ (Dissociation Constant), *K*_*i*_ (Inhibition Constant), or *IC*_50_ (Half Maximal Inhibitory Concentration). To clearly label the interacting partners among chains in PDB protein complexes, we identified the chain components involved in the direct pairwise interactions of a minimum interaction unit. This approach aligns with structure-based benchmark database for protein-protein binding affinity,^31^ which is an existing benchmark recording the chain components of pairwise binding in protein-protein complexes. Our focus was narrowed to complexes containing either two or three types of proteins, excluding more complex assemblies to facilitate the conversion of intricate protein-protein interactions into direct pairwise relationships. The labeling process followed specific patterns and corresponding rules shown in Figure 2: (i) In cases with only two chains belonging to separate proteins, the simplest scenario, we designated these two chains as the interaction components. (ii) For complexes containing more than two chains, we conducted a manual structural examination to ascertain the chains involved in direct interaction. Where structure alone was insufficient for determination, the related publications were consulted for clarification, for instance, checking the precise interaction composition measured in the experimental setup. (iii) If a complex comprised identical units, such as a symmetrical oligomeric complex, we extracted the smallest unit for further analysis.

These guidelines were adhered to in our manual labeling process, with the exception of a few exceptionally challenging cases. Ultimately, this resulted in a collection of 2,285 labeled pairwise PDB structures.

**Figure 2:**
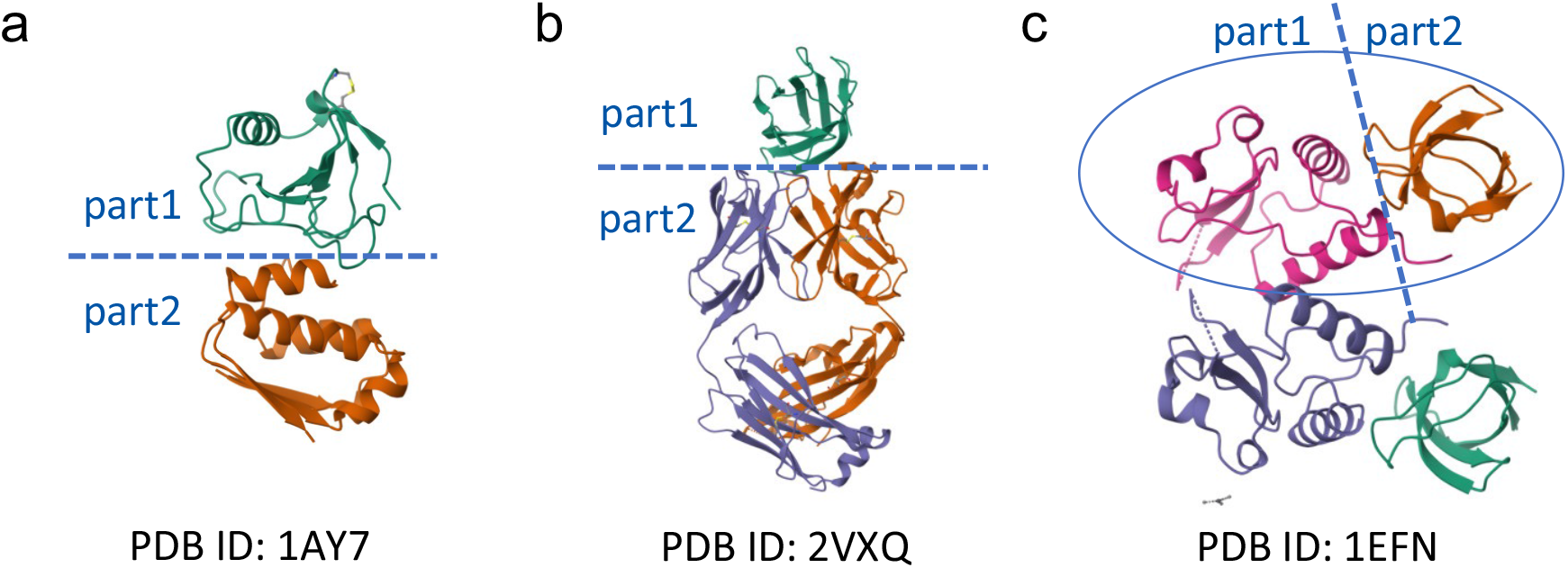
Common scenarios when dealing with protein-protein complexes. (a) The complex is composed of only 2 chains. (b) The complex is composed of multiple chains (more than 2). (c) The complex contains identical units, e.g., a symmetrical dimeric complex here.

### Data preparation

In preparing the dataset for training our ProAffinity-GNN model aimed at predicting binding affinities, we undertook a rigorous selection process. Initially, we filtered out any entries containing DNA/RNA structures due to their incompatibility with our model’s focus, resulting in a dataset comprising 2,270 entries. Each entry contains PDB files detailing the atomic-level structures of protein-protein complexes. To enhance the dataset’s utility for our analyses, we converted these PDB files into PDBQT format using AutoDockFR.^35^ This process also involved adding polar hydrogens for better molecular representation.

The dataset is characterized by binding affinity values primarily denoted by *K*_*d*_, alongside a smaller proportion labeled with *K*_*i*_ or *IC*_50_, all of which are conceptually analogous. To standardize these values and address their typically low magnitudes, we adopted the negative logarithm to base 10 (*pK*_*a*_) as the uniform measure for binding affinity, formulated as:

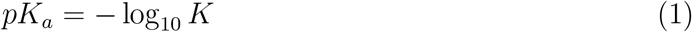

In this context, *K* can represent any of the affinity values (*K*_*d*_, *K*_*i*_, or *IC*_50_), with mol/L as the unit of measurement.

Additionally, we extracted sequence information for each protein complex from Protein Data Bank^36^ by retrieving the FASTA sequence, ensuring a comprehensive representation of each entry’s molecular structure and properties.

To maintain the integrity of our model evaluation and ensure no overlap with benchmark datasets used for testing, we removed any entries from our dataset that were present in those benchmarks. This resulted in a refined dataset of 2,133 entries. The distribution of the target value is shown in Figure 3.

**Figure 3:**
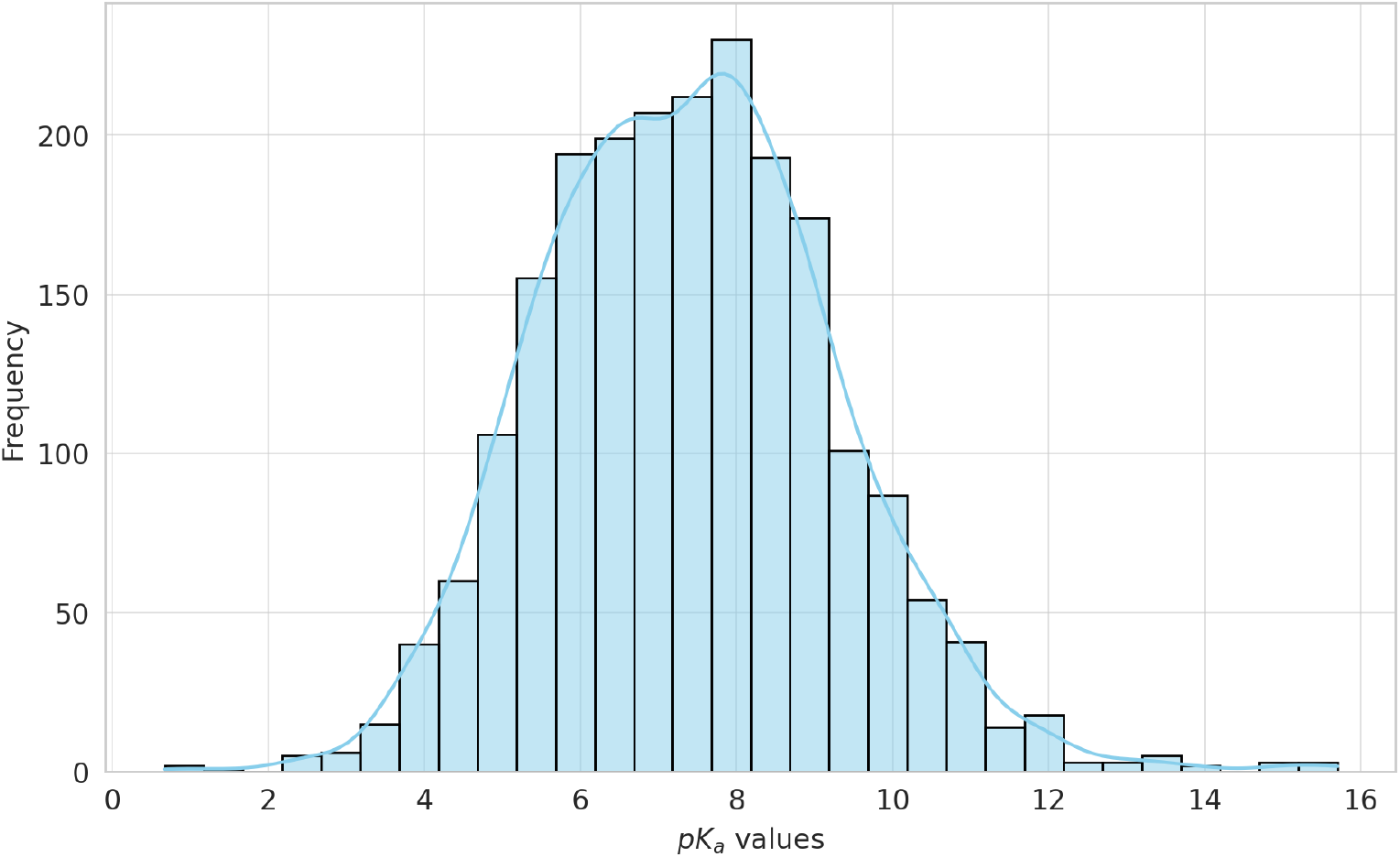
Distribution of the target values.

We subsequently divided this dataset into a training set and a validation set using an 80:20 ratio. Furthermore, we organized the data into five distinct groups, enabling a 5-fold cross-validation approach.

### Graph construction

Acknowledging the role of both inter-residue contacts and individual protein structures in protein-protein binding affinity,^21^ we crafted two distinct types of graphs: the intra-molecular graph highlights interactions within each protein, while the inter-molecular graph captures interactions between protein pairs. These residue-based graphs depict individual amino acids as nodes. The adjacency matrix for intra-molecular graphs, *A*_*intra*_, is defined such that *A*_*i,j*_ = 1 if the distance between nodes *i* and *j* is within the intra-molecular cutoff, and 0 otherwise. Similarly, for inter-molecular graphs, *A*_*inter*_, edges are established between the two components of the pairwise complex based on an inter-molecular distance threshold. We set the intra-molecular cutoff at 3.5Å, and the inter-molecular cutoff at 15Å.

For both intra- and inter-molecular graphs (*G*_*intra*_ and *G*_*inter*_), we utilized the Evolutionary Scale Modeling-2 (ESM-2)^33^ for node embedding. ESM-2, a cutting-edge transformer-based protein language model, has been trained on a vast corpus of protein sequences, drawing from the advancements in large language models within the NLP (natural language processing) domain.^37^ Specifically, we used esm2 t33 650M UR50D, which boasts 650 million parameters and produces 1280-dimensional embeddings for each residue, offering high-quality representations of protein sequences. The model’s efficiency and scale make it particularly suitable for contemporary protein research.^38,39^ To ensure comprehensive sequence representation, we input the FASTA sequences corresponding to each protein complex into the ESM-2 model, whose outputs are used as node embeddings. This approach allows us to capture the most complete sequence information available. For complexes with multiple chains, embeddings are generated for each chain individually. These embeddings are then aligned with the residues from the PDB files to establish accurate edge connections based on the three-dimensional protein structure.

Drawing on the principles of OnionNet,^40^ which was developed for predicting proteinligand binding affinities, we implemented a similar onion-like model to methodically categorize the spatial relationships of atom pairs within protein complexes. This model segregates space into concentric layers or “shells”, determined by the boundary distance *D*, aligned with our predefined cutoff. The space within *D* is divided into *N* bins or shells, each with a thickness 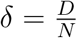, enabling us to classify atom pairs based on which shell they occupy. Atom pairs exceeding the distance *D* are considered to be in the outermost shell.

Our model accounts for seven atom types (A, C, OA, N, SA, HD, NA) found in PDBQT format files, leading to 28 distinct atom-pair types, considering the symmetry in pair designation. For each shell, we quantify the occurrence of each atom-pair type, constructing a *N ×* 28 dimensional vector that serves as the edge embedding. Specifically, we set *N* = 10, with shell thickness *δ* = 1.5Å for inter-molecular graphs and *δ* = 0.35Å for intra-molecular graphs, thereby providing a structured and detailed representation of the molecular interactions within and between protein molecules.

### Network

First, we introduced the overall architecture of the network. ProAffinity-GNN’s architecture (Figure 4) is structured to process three distinct graphs: one inter-molecular (*G*_*inter*_) and two intra-molecular (*G*_*intra*_) graphs. The network employs multiple Graph Neural Network (GNN) layers featuring an attention mechanism to distill and learn intricate features from each graph. After these GNN layers update the node features within each graph, a pooling operation aggregates these updated features across all nodes in each graph to capture comprehensive graph-level representations.

**Figure 4:**
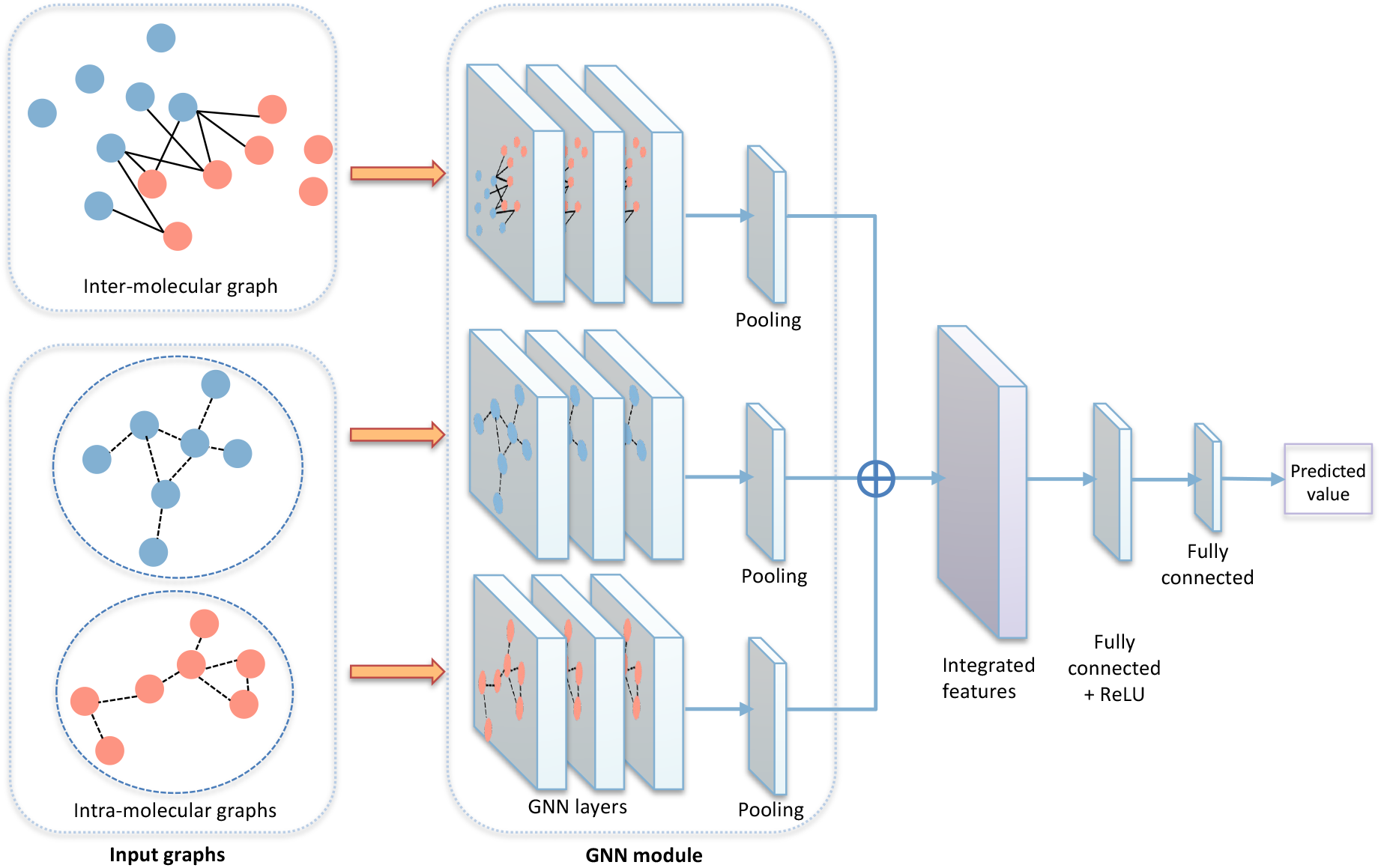
ProAffinity-GNN Architecture: Processes three distinct graphs (one intermolecular and two intra-molecular) using GNN layers with an attention mechanism, followed by pooling for graph-level feature aggregation, concatenation for comprehensive interaction insights, and fully connected layers with ReLU activation for regression output.

These aggregated features from each graph are then concatenated, ensuring that the combined feature vector encapsulates comprehensive information from both the protein-protein interaction surface and the individual protein components. The network proceeds with several fully connected layers activated by ReLU functions, with the exception of the final layer, which employs a linear transformation to produce the regression output, predicting the binding affinity.

Then, we delve into the Graph Attention Mechanism here, a pivotal component of our GNN layers, to elucidate how it dynamically influences feature learning on graph-structured data. Graph Attention Network (GAT)^41^ module is a sophisticated mechanism designed to enhance feature learning on graph-structured data. GATs distinguish themselves by allocating variable importance to neighboring nodes through attention coefficients, thus enabling dynamic feature aggregation based on the relevance of each neighbor’s information. This method is particularly advantageous in biological networks and protein-protein interaction (PPI) studies,^9,10,42^ where the significance of interactions between nodes varies.

In practice, a GAT layer updates each node’s feature vector by first computing attention coefficients (*α*_*ij*_) for every neighbor *j* of node *i*, using a shared attention mechanism. These coefficients are calculated as follows:

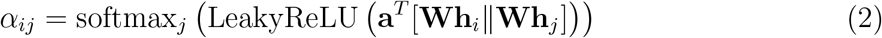

Here, **h**_*i*_ and **h**_*j*_ represent the feature vectors of nodes *i* and *j* respectively, **W** is a weight matrix applied to every node, ∥ denotes concatenation, and **a** is the attention mechanism’s weight vector. The softmax function ensures that the attention coefficients across all neighbors sum to 1, allowing for a normalized aggregation of neighbor features.

The updated feature vector for node *i* 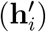 is then derived as a weighted sum of its neighbors’ transformed features, with weights determined by the attention coefficients:

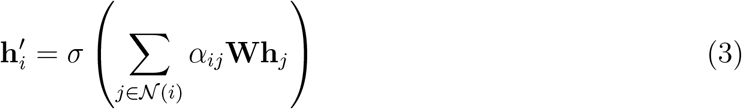

where *σ* denotes a non-linear activation function such as ReLU. This process emphasizes more informative neighbors during feature aggregation, enhancing the model’s ability to capture the complex dynamics of protein-protein interactions.

In our model, we integrate the AttentiveFP^43^ framework (Figure 5) as the core of our GNN to process the input graphs efficiently. AttentiveFP enhances the model by first individually updating the nodes within a graph. It then introduces a novel approach by creating a virtual super node that connects to all other nodes, encapsulating the entire graph’s information. This methodology allows AttentiveFP to harness both detailed local node features and broader graph-level insights, facilitating more accurate predictions for graph-based tasks.

**Figure 5:**
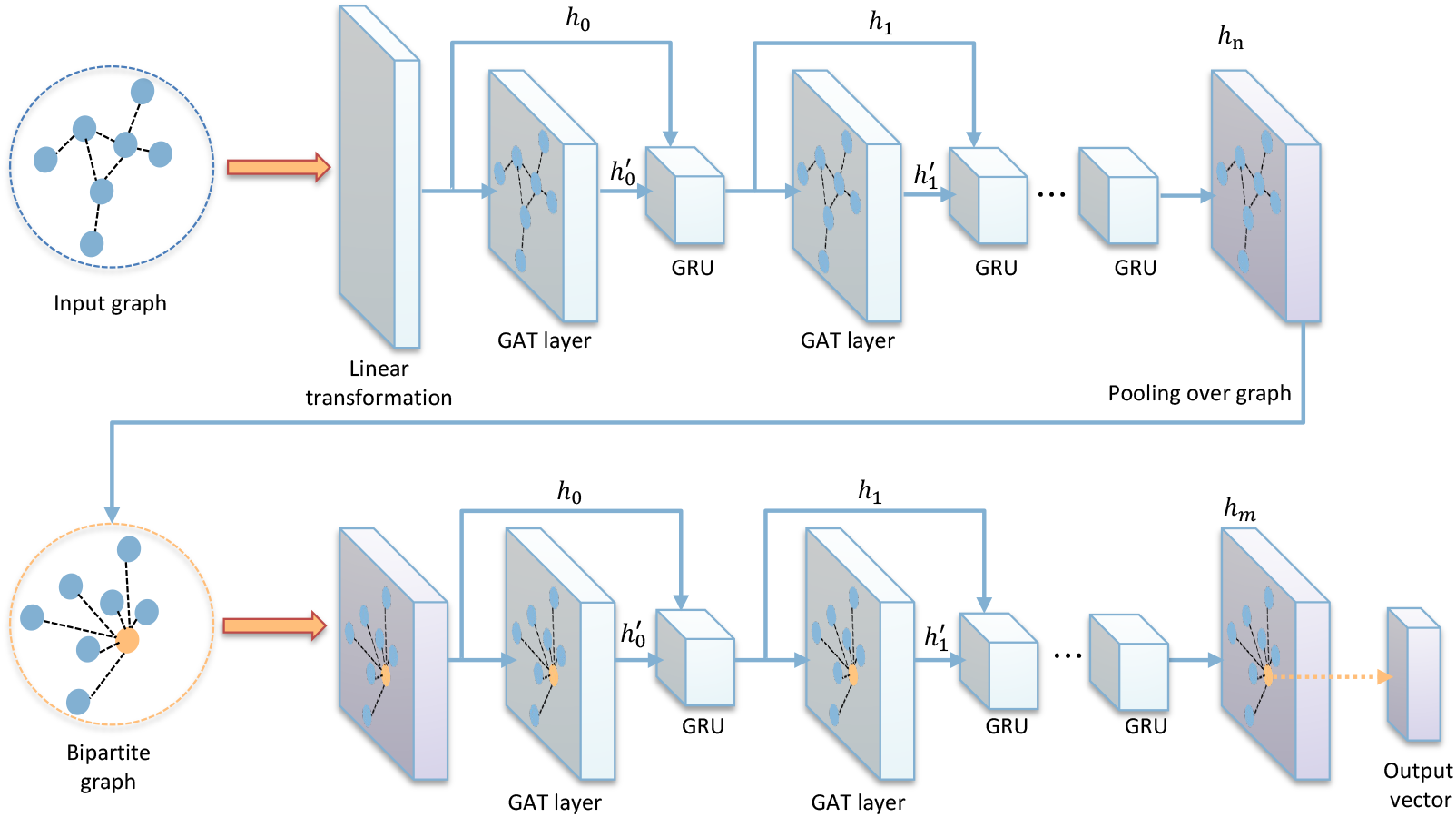
AttentiveFP Module: Begins with dimensionality adjustment via a fully connected layer, followed by feature refinement through cycles of GAT and GRU. Incorporates a super virtual node for comprehensive graph representation, leading to a bipartite structure that enhances information aggregation. Finalizes with ReLU-activated fully connected layers to distill graph-level features into a precise predictive value, merging local and global graph data for improved accuracy.

The AttentiveFP architecture operates as follows (see Figure 5): Initially, a fully connected layer adjusts the dimensionality of the node vectors to a uniform size. Subsequently, the Graph Attention Network (GAT) layer updates the initial node features, which are then processed by a Gated Recurrent Unit (GRU). The GRU combines the current input state *x*_*t*_ with the previous hidden state *h*_*t*−1_, as shown in the equation:

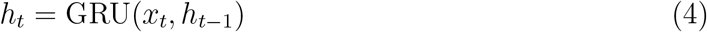

Here, *h*_*t*_ represents the output state at time *t*. Within AttentiveFP, the GRU’s inputs include node features both before and after being updated by the GAT layer, corresponding to **h**_*i*_ and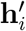 from the GAT process (Eq. 2 and Eq. 3). The details of GRU will be demonstrated in this session later.

This node updating cycle is repeated multiple times to refine the feature representations. Following the node updates, a comprehensive graph representation is formed by adding a super virtual node that symbolizes the entire graph, linking it to every individual node. This constructs a bipartite graph where the super node serves as a central hub, aggregating information across the graph. The feature vector of this super node, derived from pooling over the entire graph, undergoes further repeated updates using the same mechanism.

Finally, the model employs a series of fully connected layers activated by ReLU functions to distill the enriched graph-level features into a singular predicted value. Through this structured approach, AttentiveFP adeptly combines local and global graph information, significantly enhancing the model’s predictive accuracy for complex graph-based tasks.

The detailed operation of the GRU module for updating node features is governed by the following formulas.

- Reset Gate determines how much of the past information needs to be forgotten:

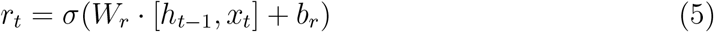

where *r*_*t*_ represents the reset gate vector at time step *t, σ* is sigmoid activation function, [*a, b*] denotes the concatenation of vector *a* and *b, h*_*t−*1_ and *x*_*t*_ indicate the previous hidden state and current input state respectively, and *W*_*r*_ and *b*_*r*_ are weight matrix and bias of the reset gate respectively.

- Update Gate, which has a similar structure to Reset Gate, determines the blend of the previous state and new input to form the current state, finely balancing memory retention and updating:

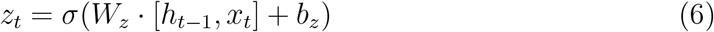

- Candidate Hidden State represents a potential new state combining the current input with relevant past information:

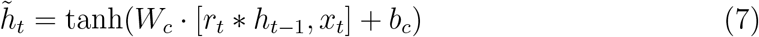

where *∗* represents element-wise multiplication.

- The final hidden state *h*_*t*_ is a blend of the past state and the candidate hidden state, as moderated by the update gate:

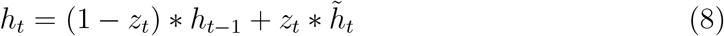

### Model training

We trained our model to predict *pK*_*a*_ values over a maximum of 25 epochs, using an Adam optimizer^44^ with a learning rate of 0.0005 and weight decay of 0.001. Training batches were set at 16, with a dropout rate of 0.5 to avoid overfitting, and early stopping was employed based on validation set performance to optimize training duration and efficiency. The objective was to minimize Mean Squared Error (MSE) Loss, conducted on an NVIDIA RTX A6000 GPU.

### Evaluation metrics

For assessing our model’s performance, we adopted Mean Absolute Error (MAE) and Pearson’s Correlation Coefficient (R) as primary metrics, aligning with standards set by existing benchmarks. These metrics are defined as follows:

MAE quantifies the average magnitude of errors between predicted (*ŷ*_*i*_) and actual (*y*_*i*_) binding values across *N* samples:

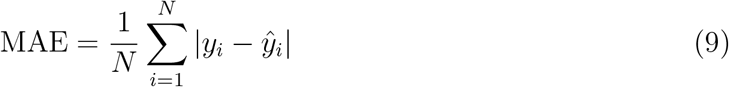

Pearson’s Correlation Coefficient (R) measures the linear correlation between the predicted and actual binding values, providing insight into the prediction accuracy and directionality:

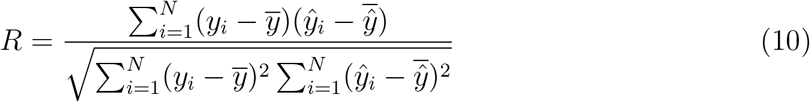

Here, 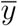 and 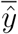 represent the mean values of actual and predicted binding values, respectively.

To facilitate comparison with prior work, ^28^ we converted our model’s output from *pK*_*a*_ to Δ*G* (change in Gibbs free energy upon binding), measured in kcal/mol, using the relationship between Δ*G* and *K* (*K*_*d*_, *K*_*i*_, or *IC*_50_):

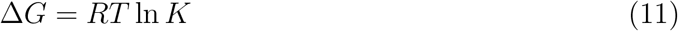

Given *R* (universal gas constant) as 1.987 *×* 10^*−*1^ and *T* (temperature) as 298, the conversion from *pK*_*a*_ to Δ*G* is straightforward due to their linear relationship:

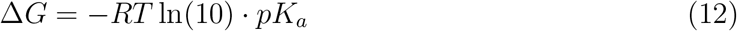

Although our model targets *pK*_*a*_, converting to Δ*G* allows direct comparison with existing benchmarks. For MAE, this entails a simple linear scaling, while *R* remains unaffected by the conversion. The following section presents results in the transformed Δ*G* format for consistency with established benchmarks.

## Results and discussion

We conducted a detailed assessment of the ProAffinity-GNN model, starting with 5-fold cross-validation on our training set. Following this, we compared the model’s performance against established methods using specialized test sets for protein-protein binding affinity prediction. To further validate our framework’s design, we also carried out an ablation study, which helped identify the impact of specific components on the overall framework efficacy.

### Performance through 5-Fold cross-validation on the training set

To ascertain the generalizability and robustness of our model, we employed 5-fold crossvalidation on a training dataset comprising 2,133 entries. This method involved segmenting the dataset into five equal parts, cyclically using one as the validation set while the remainder served for training. This cycle was repeated five times, ensuring each subset was used for validation once. The model’s performance across these folds was recorded and averaged, with the standard deviation provided. Our model demonstrated a mean correlation coefficient *R* = 0.629 *±* 0.056 and a mean absolute error (MAE) of MAE = 1.62 *±* 0.08 kcal/mol, showing a notable improvement over the previous PPI-Affinity model,^28^ which reported an *R* value of 0.53 using an ensemble model on their development set.

### Comparative analysis with existing methods on benchmark sets

To rigorously assess the performance of our ProAffinity-GNN model, we compared it against existing methods using a series of established benchmark datasets for protein-protein binding affinity prediction. The first test set comprises 79 data points, extracted from “structurebased benchmark database for protein-protein binding affinity”,^31^ which includes protein-protein complexes that may consist of two or more protein chains. The second test set, developed in the context of the PPI-Affinity project, is drawn from the PDBbind database and features 90 data points representing complexes solely composed of two protein chains. While this set includes one homo-dimer complex, despite homo-dimers not being considered in our training data, we opted to retain it for completeness. To provide a holistic view, we also combined these data from both test sets, yielding a comprehensive dataset for evaluation. The results of these analyses are presented in Table 1, highlighting the comparative performance of ProAffinity-GNN against the existing methods.

**Table 1:**
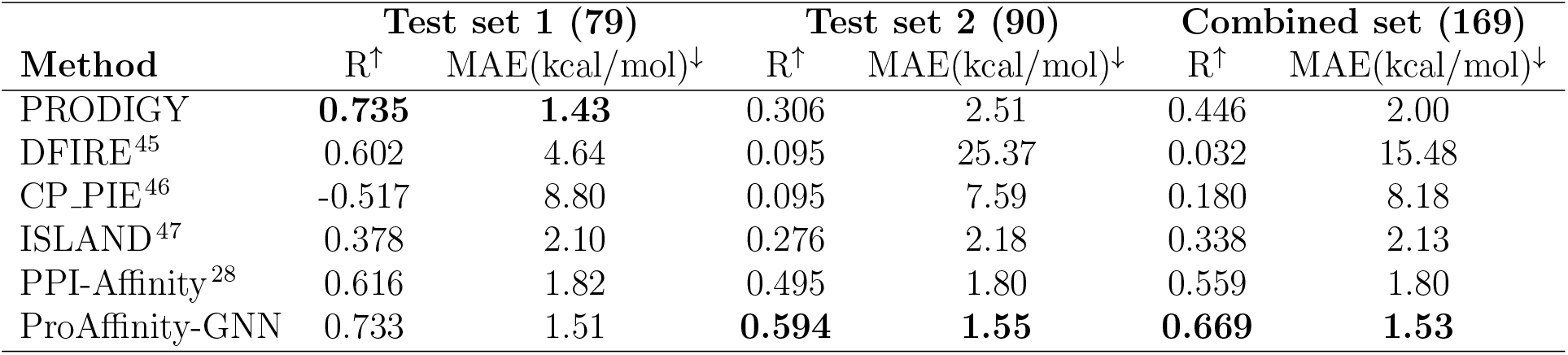
Performance compared with existing methods on benchmarks.

Our ProAffinity-GNN model exhibits performance on par with PRODIGY on test set and demonstrates superior stability and accuracy across the remaining datasets, particularly test set 2 and the combined set. This suggests that our model offers notably enhanced generalizability compared to prior methods, which often show significant performance degradation on more challenging datasets.

### Evaluation on another external dataset: SKEMPI

To substantiate the effectiveness of our approach, we conducted an assessment of our model using a segment of the SKEMPI (Structural database of Kinetics and Energetics of Mutant Protein Interactions) 2.0 database.^48^ This database is dedicated to exploring the dynamics of mutant protein interactions, with a particular emphasis on how mutations influence protein-protein binding affinity. It is a valuable resource for benchmarking in the realm of binding affinity prediction.^27,49,50^ The raw database encompasses 345 wild-type protein structures and an expansive collection of 7,085 mutants, each annotated with binding affinity in *K*_*d*_. We took a subset previously established, consisting of 26 wild-type and 151 mutant structures. ^28^ We eliminated duplicate structures, resulting in a final compilation of 26 wild-type and 140 mutant protein-protein complexes. Utilizing the EvoEF2 toolkit, ^51^ we generated complete structures for the mutant complexes. Then we evaluated ProAffinity-GNN’s performance compared with PPI-Affinity, as detailed in Table 2.

**Table 2:** Performance comparison on SKEMPI datasets.

| Method | SKEMPI (166) |  | SKEMPI mut (140) |  |
| --- | --- | --- | --- | --- |
|  | R<sup>†</sup> | MAE(kcal/mol)<sup>↓</sup> | R<sup>†</sup> | MAE(kcal/mol)<sup>↓</sup> |
| PPI-Affinity | 0.773 | 1.33 | 0.770 | 1.37 |
| ProAffinity-GNN | **0.811** | **1.10** | **0.826** | **1.08** |

Notably, our model was not specifically trained on protein-protein complexes involving mutations, yet it still generated a desirable result and managed to outperform the existing model on datasets comprising mutant data. This accomplishment underscores ProAffinity-GNN’s robustness and its capacity to generalize across a diverse range of protein interactions, including those not represented in the training dataset. The model’s strong performance on mutation-affected binding affinities highlights its potential broad applicability such as further protein function research and therapeutic development targeting protein interactions.

### Ablation study

To ascertain the efficacy of the ProAffinity-GNN framework’s design, we conducted an ablation study. This study aimed to dissect the impact of the important components of the GNN structure on the model’s performance, thereby clarifying their individual contributions to the framework’s success in predicting protein-protein binding affinity.

Our ablation study comprised several GNN model variants, each with a specific modification—either through the removal or alteration of a component. The baseline model integrates an AttentiveFP approach with a combination of three graphs: one inter-molecular graph and two intra-molecular graphs. The variants tested include:

- **Variant A**: Utilizes a Graph Attention Network (GAT) baseline with three layers and includes edge features, accompanied by three graphs.
- **Variant B**: Features an AttentiveFP baseline combined with an inter-molecular graph.
- **Variant C**: Incorporates an AttentiveFP baseline with two intra-molecular graphs.

To maintain experimental consistency, each variant underwent evaluation using the same training procedure, dataset, and metrics as the standard model. The outcomes of our ablation study are shown in Table 3 and Figure 6. From the results, we observe the following:

1. Solely employing GAT layers within the GNN model results in a performance decline. This indicates that GATs alone may not be optimally suited for this task. Conversely, the AttentiveFP structure more effectively aggregates information across the comprehensive graph.
2. A model that encompasses the entire complex, including both inter-molecular contacts and individual protein details, achieves superior results compared to models that focus on either aspect in isolation.
3. Respectable results can be achieved using individual protein structures, even in the absence of explicit inter-molecular interaction features. Variant B, which includes an AttentiveFP baseline augmented with an inter-molecular graph, incorporates edge features that represent detailed interactions between protein groups based on 3D structural data. Variant C, lacking an inter-molecular graph, does not explicitly account for these interactions, relying instead on two separate intra-molecular graphs. De-spite this, the performance of Variant C is comparable to Variant B, suggesting that the model can also capture essential information from individual protein structures without explicit inter-molecular graph features.

**Table 3:** Detailed results of the ablation study.

| Method/Test | 5-fold CV |  | Test set 1 (79) |  | Test set 2 (90) |  | SKEMPI subset |  |
| --- | --- | --- | --- | --- | --- | --- | --- | --- |
| | $R^\dagger$ | MAE(kcal/mol)$^\downarrow$ | $R^\dagger$ | MAE(kcal/mol)$^\downarrow$ | $R^\dagger$ | MAE(kcal/mol)$^\downarrow$ | $R^\dagger$ | MAE(kcal/mol)$^\downarrow$ |
| Standard | 0.629±0.056 | 1.62±0.08 | 0.733 | 1.51 | 0.594 | 1.55 | 0.811 | 1.10 |
| Variant A | 0.583±0.060 | 1.65±0.07 | 0.643 | 1.63 | 0.565 | 1.70 | 0.658 | 1.50 |
| Variant B | 0.600±0.029 | 1.85±0.12 | 0.705 | 1.69 | 0.563 | 1.67 | 0.787 | 1.23 |
| Variant C | 0.610±0.062 | 1.68±0.11 | 0.650 | 1.72 | 0.586 | 1.66 | 0.790 | 1.67 |

**Figure 6:**
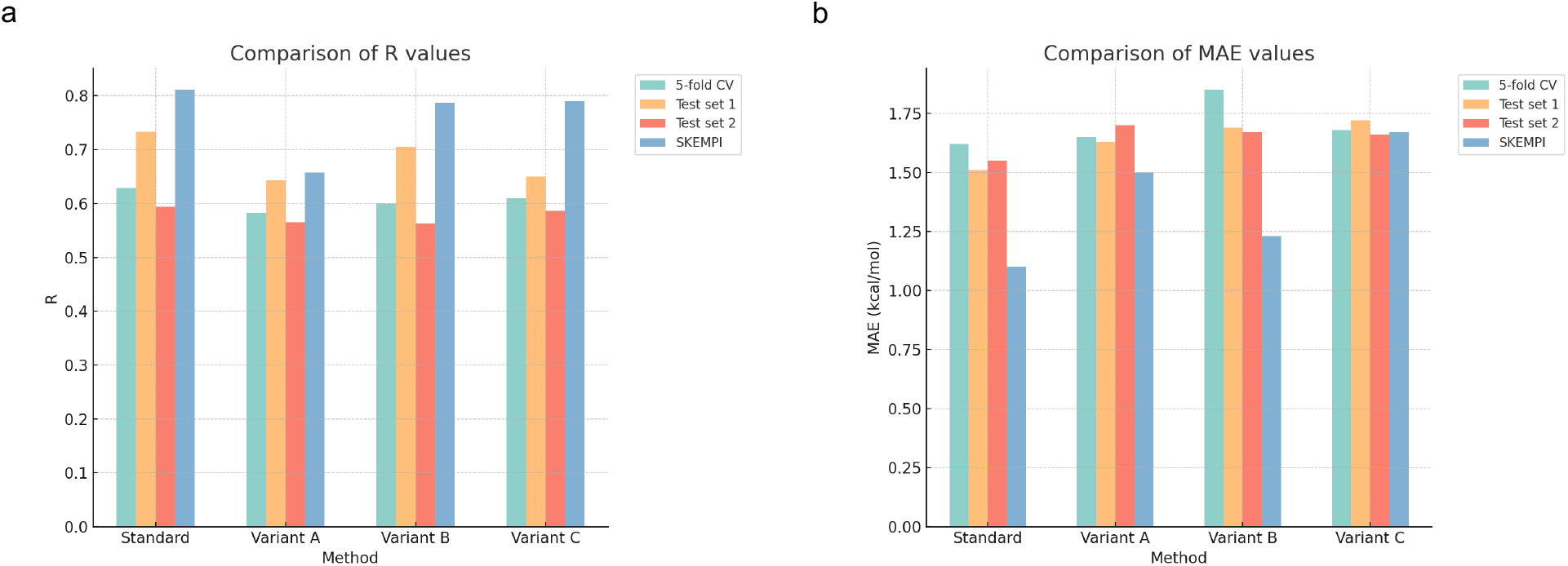
The results of the ablation study. (a) The test results of different methods with respect to R. (b) The test results of different methods with respect to MAE.

## Conclusion

The prediction of protein-protein binding affinity occupies a pivotal role in the study of protein-protein interactions, heralding advancements in protein engineering and drug discovery. Despite its significance, the field is hampered by a notable scarcity of effective and efficient methodologies. A critical challenge is the significant lack of comprehensive data. Moreover, the processing procedures of existing methods are often overly complex and tailored to only a subset of the relevant concerns. In response to these limitations, we have developed an enriched dataset based on PDBbind, focusing on pairwise interacting protein-protein complexes, and have manually added labels for identifying interacting two groups, each of which may contain multiple chains based on experimental setups. We introduce a novel deep learning approach, ProAffinity-GNN, which leverages a protein language model and graph neural networks, integrating the intuitive spatial structure with the protein sequence that holds a lot of potential messages. This method is also distinguished by its simplicity in processing complexes without necessitating the cumbersome computation of physicochemical properties. Extensive evaluations demonstrate that ProAffinity-GNN not only achieves superior performance compared to existing techniques but also exhibits remarkable generalization capabilities across diverse external datasets, yielding more consistent prediction outcomes.

The journey towards refining protein-protein binding affinity prediction is far from over, with significant strides still to be made in dataset development and model innovation. A pressing issue remains the absence of a universally accepted evaluation dataset, which is crucial for the equitable validation of model performance. It is our hope that the field will witness increased research efforts aimed at overcoming these challenges, further propelling the advancement of this critical area of study.

## Acknowledgement

This research was supported by Singapore Ministry of Education (MOE) Tier 1 RG97/22. Computations were mainly performed using the resources of the National Supercomputing Centre, Singapore (https://www.nscc.sg) and the HADLEY high-performance computing cluster of SCELSE. SCELSE is funded by Singapore’s National Research Foundation, the Ministry of Education, NTU, and the National University of Singapore (NUS), and is hosted by NTU in partnership with NUS.

